# Hepatic FoxOs induce apolipoprotein M and are required for sphingosine-1-phosphate to bind high density lipoproteins

**DOI:** 10.1101/804336

**Authors:** María Concepción Izquierdo, Niroshan Shanmugarajah, Samuel X. Lee, Rebecca A. Haeusler

## Abstract

The FoxO family of transcription factors play an important role in mediating insulin action on glucose, lipid, and lipoprotein metabolism. Liver-specific triple FoxO knockout mice (L-FoxO1,3,4) have defects in expression of genes related to glucose production, bile acid synthesis, and high density lipoprotein (HDL)-cholesterol uptake. We have now identified Apolipoprotein M (Apom) as a novel transcriptional target of liver FoxO. ApoM is a liver-secreted apolipoprotein that is bound to HDL in the circulation, and it serves as a chaperone for the bioactive lipid, sphingosine-1-phosphate (S1P). Several recent studies have demonstrated that S1P bound to ApoM induces unique effects, compared to S1P bound to albumin. We now show that liver FoxOs are required for ApoM mRNA and protein expression, and that ApoM is a transcriptional target of FoxOs. Moreover, while total plasma S1P levels are similar between control and L-FoxO1,3,4 mice, S1P is nearly absent from HDL in L-FoxO1,3,4 mice, and is instead increased in the lipoprotein depleted fraction. We also observed that leptin receptor deficient db/db mice have low hepatic Apom mRNA, and low levels of ApoM and S1P in HDL, without changes in total plasma S1P. These data demonstrate that FoxO transcription factors are novel regulators of the ApoM-S1P pathway, and indicate a potential link between hepatic insulin action and HDL function.

## INTRODUCTION

Insulin resistance is a major driver of dyslipidemia and cardiovascular risk. However, the mechanisms linking alterations in insulin signaling with dyslipidemia are incompletely understood. The liver is a key tissue for integrating the signaling of insulin and glucose: it is responsible for maintaining blood glucose concentration, it acts as a reservoir for delivery of glucose to the bloodstream when it is necessary^1^, and it is a key target for insulin’s inhibition of hepatic glucose production^2^. Moreover, the liver is critical to the regulation of lipoproteins: very low-density lipoprotein (VLDL) particles and nascent high-density lipoprotein (HDL) particles are synthesized and secreted into the circulation by the liver. Hepatocytes are the primary site of uptake for LDL particles, and HDL particles also return to the liver to deliver cholesterol removed from other tissues through the process of reverse cholesterol transport^3^. Therefore, liver insulin signaling is a potential contributor to insulin resistance-associated dyslipidemia.

FoxO transcription factors are critical mediators of insulin action on gene expression in hepatocytes, where they are involved in regulating glucose and lipid metabolism^4–8^. Consistent with this, mice with liver-specific knockout of all three insulin-sensitive FoxO isoforms (L-FoxO1,3,4) have reduced hepatic glucose production and increased de novo lipogenesis^5, 9, 10^. These phenotypes are associated with altered expression of genes related to glucose versus fatty acid production, including reduced glucose-6-phosphatase catalytic subunit (*G6pc*) and increased glucokinase^5, 9, 11, 12^. L-FoxO1,3,4 mice also have defects in bile acid synthesis and HDL-mediated reverse cholesterol transport^13, 14^.

Based on unbiased transcriptomics from livers of L-FoxO1,3,4 mice^5^, one of the most downregulated genes in the livers of these mice is Apolipoprotein M (*Apom*). ApoM is a secreted protein that is bound to lipoprotein particles, and is predominantly enriched–more than 95%–in HDL^15^. ApoM is a chaperone for sphingosine-1-phosphate (S1P) in plasma HDL^16^. S1P is a bioactive sphingolipid that signals through a series of G protein-coupled receptors (S1P receptors 1-5) present on a variety of cell types^17, 18^. In plasma, around 65% of S1P is carried by HDL-bound ApoM and the remainder is found in the lipoprotein depleted (LPD) fraction, presumably associated to albumin^16, 19^. Importantly, S1P induces differential effects depending on whether it is associated with ApoM or albumin^20, 21^. This is exemplified by the ability of the HDL-ApoM-S1P complex to enhance endothelial function^16, 22–24^.

In this work, we report that hepatic FoxOs promote hepatic ApoM expression and are required for S1P to associate with HDL particles. This suggests a novel mechanism whereby hepatic FoxOs determine HDL composition and function.

## RESULTS

### FoxOs are required for hepatic ApoM expression

To examine the effects of hepatic FoxO deletion on apolipoprotein gene expression, we queried our microarrays from prior experiments^5^. We found that livers from L-FoxO1,3,4 mice had a substantial reduction in the mRNA expression of *Apom*, compared to littermate controls. To confirm this, we carried out qPCR and western blots from liver tissue and found that L-FoxO1,3,4 mice have ∼90% reductions in *Apom* mRNA expression and nearly undetectable ApoM protein in liver (Fig. 1a-b).

**Figure 1.**
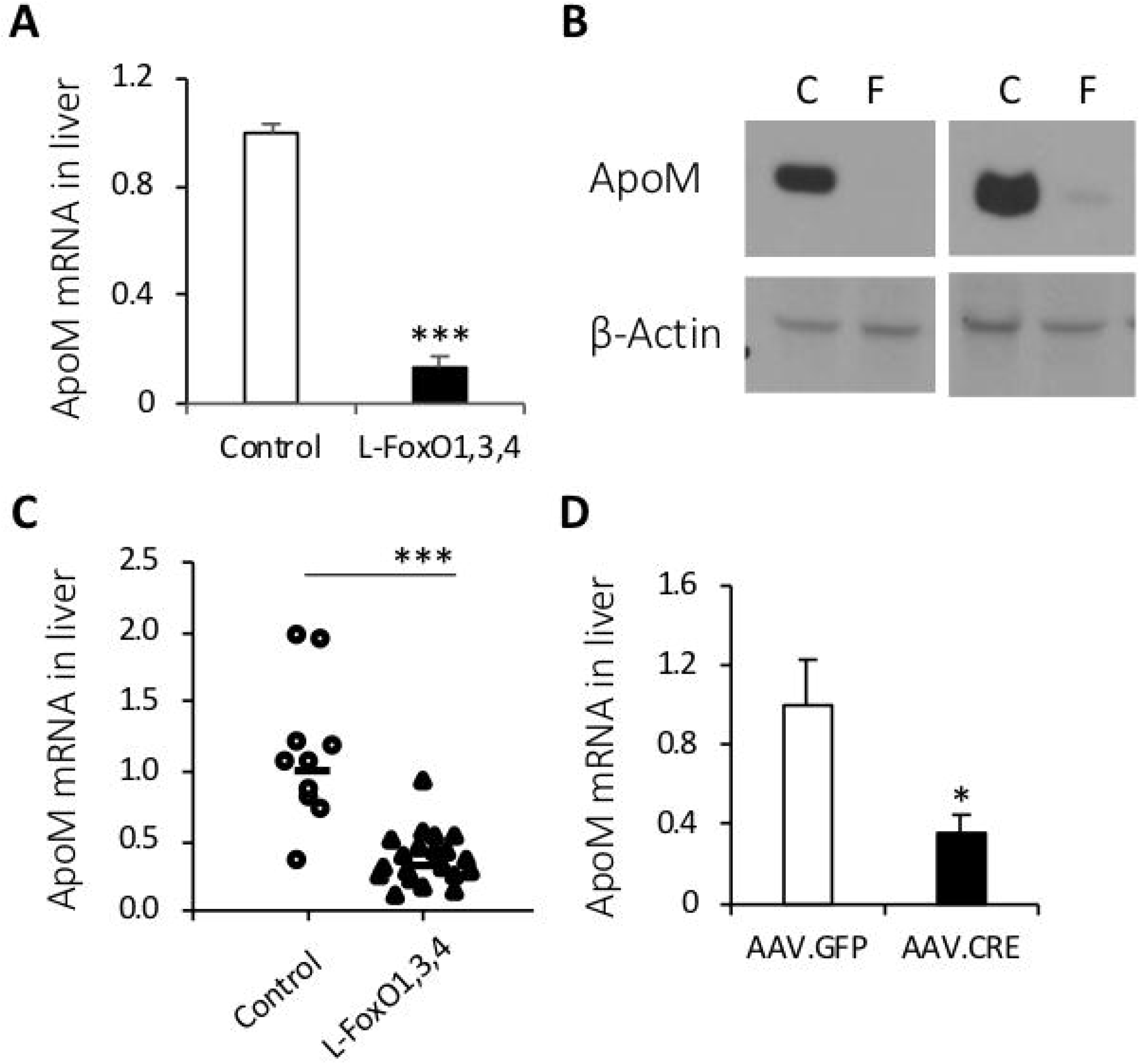
ApoM expression in liver. **A)** Hepatic *Apom* gene expression in adult male mice (n=5-8/group). **B)** Representative western blots of ApoM in liver lysates. **C)** Hepatic *Apom* gene expression from mice of both sexes sacrificed on postnatal day 2 (n=10-20/group). **D)** Hepatic *Apom* gene expression from acute knockdown via AAV8.Tbg.Cre in adult male *Foxo1^fl/fl^*, *Foxo3^fl/fl^*, and *Foxo4^fl/Y^* mice (*n* = 5/group). Values are shown relative to littermate controls. Data are presented as mean ± SEM. *p<0.05, ***P<0.001 by student’s t-tests.

The effect of FoxO deletion to reduce hepatic *Apom* expression might be due to a primary effect of FoxO ablation or an acquired or compensatory defect, due to long-term genetic loss of FoxOs. Thus, we explored the hepatic *Apom* expression in neonatal mice. The decreases in *Apom* were identified as early as post-natal day 2 in neonatal L-FoxO1,3,4 mice, compared to littermate controls (Fig. 1c). To test the effect in adult mice, we examined mice with an acute depletion of FoxOs. We transduced adult FoxO1^F/F^, FoxO3^F/F^, and FoxO4^F/Y^ control mice with an adeno-associated virus expressing Cre recombinase under the hepatocyte-specific Tbg promoter (AAV8.Tbg.Cre). We verified that one month after the injection, there was a >80% depletion of *FoxO* mRNA expression as well as low levels of *G6pc*, a known target of FoxO transcriptional activation^14^. In these FoxO-depleted mice, we found that the *Apom* mRNA expression is significantly decreased (Fig. 1d). These findings demonstrate that hepatic FoxOs are required for hepatic *Apom* expression.

### ApoM is a transcriptional target of FoxO

Hepatic FoxOs have been suggested to modulate some liver metabolic pathways via indirect effects on non-hepatic tissues^25, 26^. Therefore, we examined whether FoxOs regulate *Apom* in primary hepatocytes, using mutant versions of FoxO1. The FoxO1 protein structure contains three Akt phosphorylation sites that mediate its nuclear exclusion, a transactivation domain, and a DNA binding domain (Fig. 2a)^27, 28^. We isolated primary hepatocytes from wild-type mice and transduced them with a FoxO1-ADA mutant, which has the three Akt phosphorylation sites mutated, causing FoxO1 to be constitutively nuclear^29^. The FoxO1 ADA mutant increased *G6Pc* and *Apom* expression (Fig. 2a). FoxOs can regulate gene expression by direct DNA binding or by acting as a transcriptional coregulator^30–33^. Next we evaluated if the DNA binding domain of FoxO1 is required for the induction of *Apom*. We used a FoxO1-ADA-DBD mutant, which has the ADA mutation, and it has a second mutation in the DNA binding domain. We observed that the ADA-DBD mutant of FoxO1 is unable to activate *G6Pc* or *Apom* expression (Fig. 2b). We also transduced primary hepatocytes with a dominant negative version of FoxO1 that lacks the transactivation domain (FoxO1-Delta256)^34^. We observed that FoxO1-Delta256 decreases *G6Pc* and *Apom* expression (Fig. 2c). These data show that FoxOs promote hepatic *Apom* expression by direct actions in hepatocytes, and the DNA binding and transactivation domains are required.

**Figure 2.**
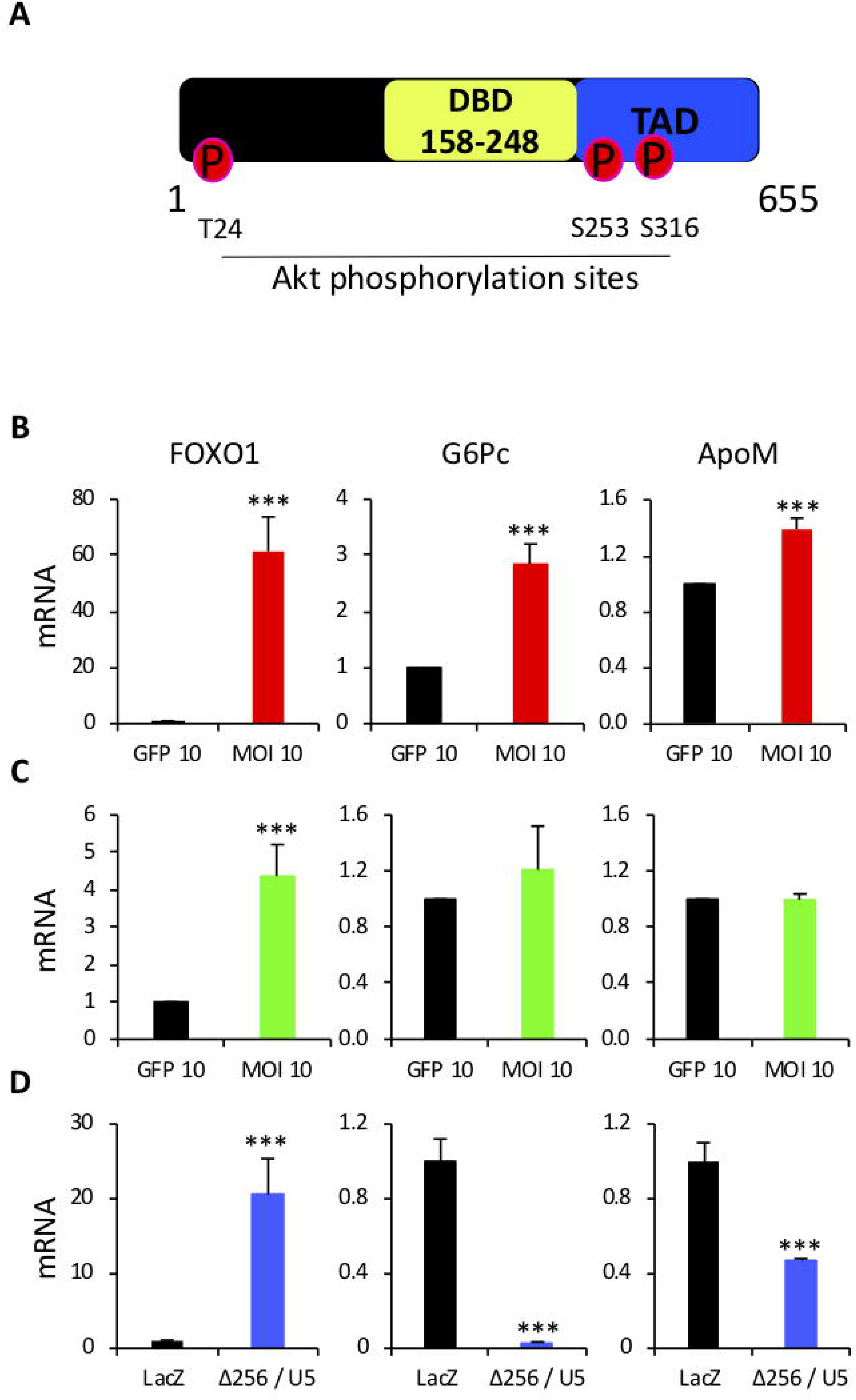
ApoM is a transcriptional target of FoxO. **A)** Schematic representation of FoxO1 protein. **B-D)** *FoxO1*, *G6Pc*, and *Apom* gene expression in primary hepatocytes from wild-type mice that were transduced with different FoxO1 mutants. **B)** FoxO1-ADA mutant: the three Akt phosphorylation sites are mutated, causing FoxO1 to be constitutively nuclear. **C)** FoxO1-ADA-DBD mutant: contains the ADA mutation and a mutation in the DNA binding domain. **D)** FoxO1-Delta256 mutant: a dominant negative version of FoxO1 that lacks the transactivation domain. Mean ± SEM of triplicates of three independent experiments. ***p<0.001 by student’s t-tests.

### FoxOs are required for S1P binding to HDL

ApoM is a secreted protein that is found mostly on HDL^16^. Thus, we examined whether FoxOs regulate the levels of ApoM in total plasma. Western blot from total plasma showed that ApoM protein levels are nearly absent in L-FoxO1,3,4 (Fig. 3a). ApoM is a chaperone of S1P^16^. We measured S1P by Liquid Chromatography-Mass Spectrometry. There were no differences in total plasma S1P levels between genotypes (Fig. 3b). Next, we examined whether ApoM and S1P are affected in plasma lipoproteins from L-FoxO1,3,4 mice. We fractionated lipoproteins by size, using fast protein liquid chromatography, and measured S1P and apolipoprotein content. This revealed that control mice have two peaks of S1P: one in the HDL, and one in the lipoprotein-depleted fractions, presumably bound to albumin (Fig. 3c). On the other hand, L-FoxO1,3,4 mice showed a marked reduction of S1P in HDL, and an increase of S1P in the lipoprotein-depleted fraction (Fig. 3c). Western blots of these fractions showed that ApoA1, the main apolipoprotein of HDL, was present in both genotypes. However, ApoM was nearly absent from L-FoxO1,3,4 mice (Fig. 3d). Therefore, hepatic FoxOs are required for ApoM and S1P association with HDL.

**Figure 3.**
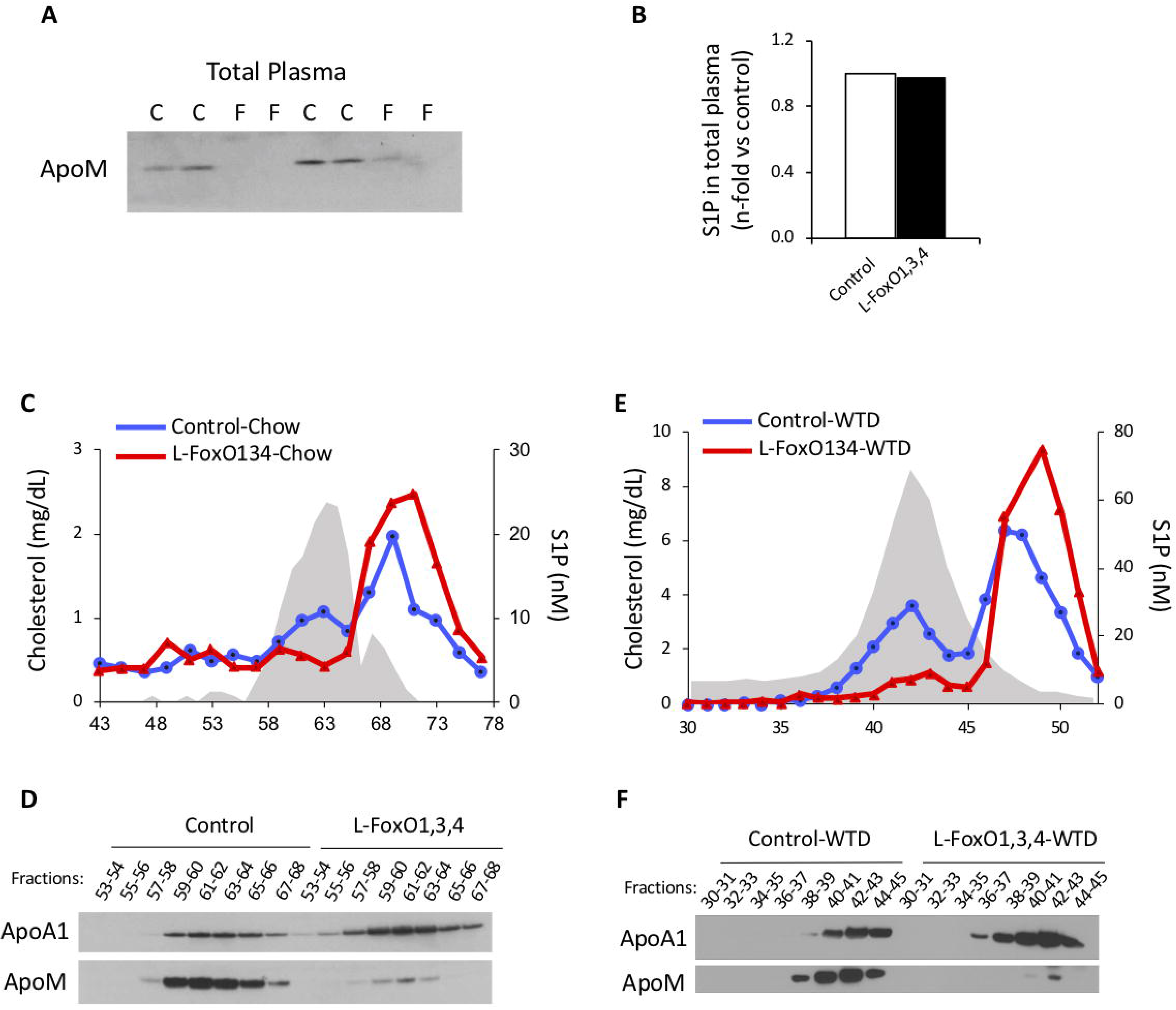
ApoM expression and S1P levels in plasma. **A)** Representative western blot of ApoM in total plasma from chow-fed mice. F=L-FoxO1,3,4 mice, C=littermate control mice (n=4/group). **B)** Total S1P levels in plasma from chow-fed mice. **C)** Distribution of S1P in plasma fractionated by size-exclusion chromatography. Cholesterol levels from the control mice are shown as a reference to demonstrate the fractions where HDL particles elute (grey color). Control mice in blue. L-FoxO1,3,4 mice in red. **D)** Western blot of ApoA1 and ApoM in lipoprotein fractions from chow-fed mice. **E)** Distribution of S1P in lipoproteins fractions from mice fed a western type diet (WTD) for three weeks. Cholesterol levels from the control mice are shown as a reference to demonstrate the fractions where HDL particles elute (grey color). Control mice in blue. L-FoxO1,3,4 mice in red. **F)** Western blot of ApoA1 and ApoM in lipoprotein fractions from western diet-fed L-FoxO1,3,4 mice.

We carried out the same analysis in mice that were fed the western diet for three weeks. Again, we observed that in the absence of hepatic FoxOs, levels of S1P and ApoM in HDL were largely depleted (Fig. 3e-f).

### Rescuing the expression of ApoM in L-FoxO1,3,4 deficient mice normalizes the S1P distribution

We next tested whether ApoM rescue in the livers of FoxO deficient mice is sufficient to normalize S1P distribution. We transduced L-FoxO1,3,4 and control mice with either an adenovirus expressing ApoM (Ad-ApoM), or a control virus (Ad-GFP). Eight days after virus injection, we harvested tissues and found that at our dose of the virus, the L-FoxO1,3,4+Ad-ApoM mice express similar levels of hepatic *Apom* mRNA as control mice transduced with Ad-GFP (Fig. 4a), demonstrating efficient rescue of *Apom*. We also measured hepatic FoxO1 expression levels and verified that there is no effect of ApoM virus injection on FoxO1 expression (Fig. 4b). We fractionated lipoproteins by sequential density ultracentrifugation, and by western blot of the HDL fractions, we confirmed the rescue of ApoM protein levels in the L-FoxO1,3,4 mice (Fig. 4c). Although there was no significant effect on total plasma S1P levels (Fig. 4d), rescuing ApoM in L-FoxO1,3,4 mice caused a normalization of the S1P distribution (Fig. 4e). We conclude that loss of ApoM is the cause of the impaired S1P association with HDL in L-FoxO1,3,4 mice.

**Figure 4.**
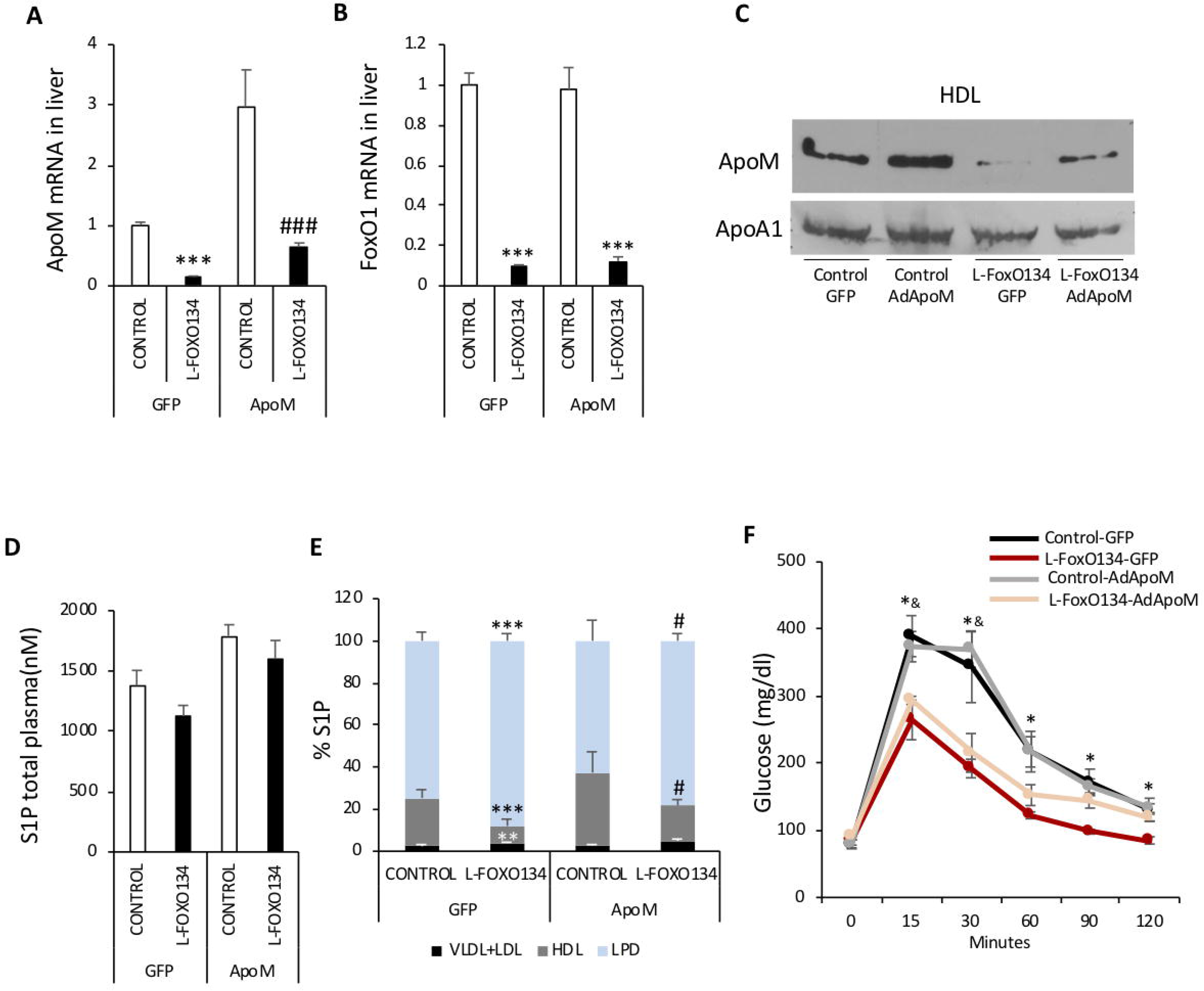
Rescue of ApoM expression. **A)** *Apom* gene expression in liver (n=4-8/group). **B)** *FoxO1* gene expression in liver (n=4-8/group). **C)** Western blot of ApoM and ApoA1 from HDL fractionated by ultracentrifugation. **D)** Total plasma S1P levels (n=3-8/group). **E)** Distribution of S1P in plasma fractions. LPD=lipoprotein depleted (n=3-8/group) **F)** Intraperitoneal glucose tolerance test (n=4-6/group) Data are presented as mean ± SEM.**P<0.01, ***P<0.001 vs controls, #P<0.05, ###P<0.001 vs L-FoxO1,3,4-GFP. In panel F, *P<0.05 control-GFP vs L-FoxO1,3,4-GFP and ^&^P<0.05 control-AdApoM vs L-FoxO1,3,4-AdApoM.

### ApoM is not involved in glucose regulation by FoxOs

Kurano *et al*.^35^ overexpressed ApoM by several-fold in wild-type mice using adenovirus gene transfer and observed that ApoM overexpression increased glucose tolerance, potentially due to increased insulin secretion. Because FoxOs are known to regulate glucose homeostasis, we tested whether rescuing ApoM in L-FoxO1,3,4 mice had any effect on glucose tolerance. As expected, we found that the L-FoxO1,3,4 mice have better glucose tolerance than littermate control mice^5, 9^, but rescuing ApoM in these mice had no effect (Fig. 4f). We conclude that ApoM is not involved in the effects of FoxOs on glucose homeostasis.

### ApoM and HDL-associated S1P are decreased in in db/db, but not diet-induced obese mice

FoxO-deficient mice are a model of constitutive insulin action onto FoxO, whose activity is normally suppressed by insulin. It’s widely held that FoxOs are constitutively active in insulin resistance, due to impaired Akt-mediated phosphorylation^36^. On the other hand, it’s also been suggested that the absence of FoxOs may mimic certain conditions of hyperinsulinemia^5, 37^ because FoxOs are exquisitely sensitive to very low levels of insulin^38^. Thus, we investigated whether the ApoM expression and S1P distribution are affected in independent mouse models of hyperinsulinemia and insulin resistance.

Leptin receptor deficient Db/db mice are a well-established model of diabetes and obesity. We examined db/db mice at two different ages, 13 and 25 weeks. The body weight, glucose, and insulin levels were higher in the db/db mice as expected (Supplementary Fig. 1a-c). We predicted that *Apom* liver expression would be increased in db/db mice, due to insulin resistance and FoxO activation. In contrast to our expectations, we observed that hepatic *Apom* expression is reduced in db/db mice (Fig. 5a-b). Consistent with this, ApoM protein levels are reduced in total plasma (Fig. 5c) and specifically in the HDL fraction from db/db mice (Fig. 5d-e). While there were no differences in total plasma S1P levels (Fig. 5f), the db/db mice showed a reduction of S1P in HDL, and an increase of S1P in the lipoprotein depleted fraction (Fig. 5g).

**Figure 5.**
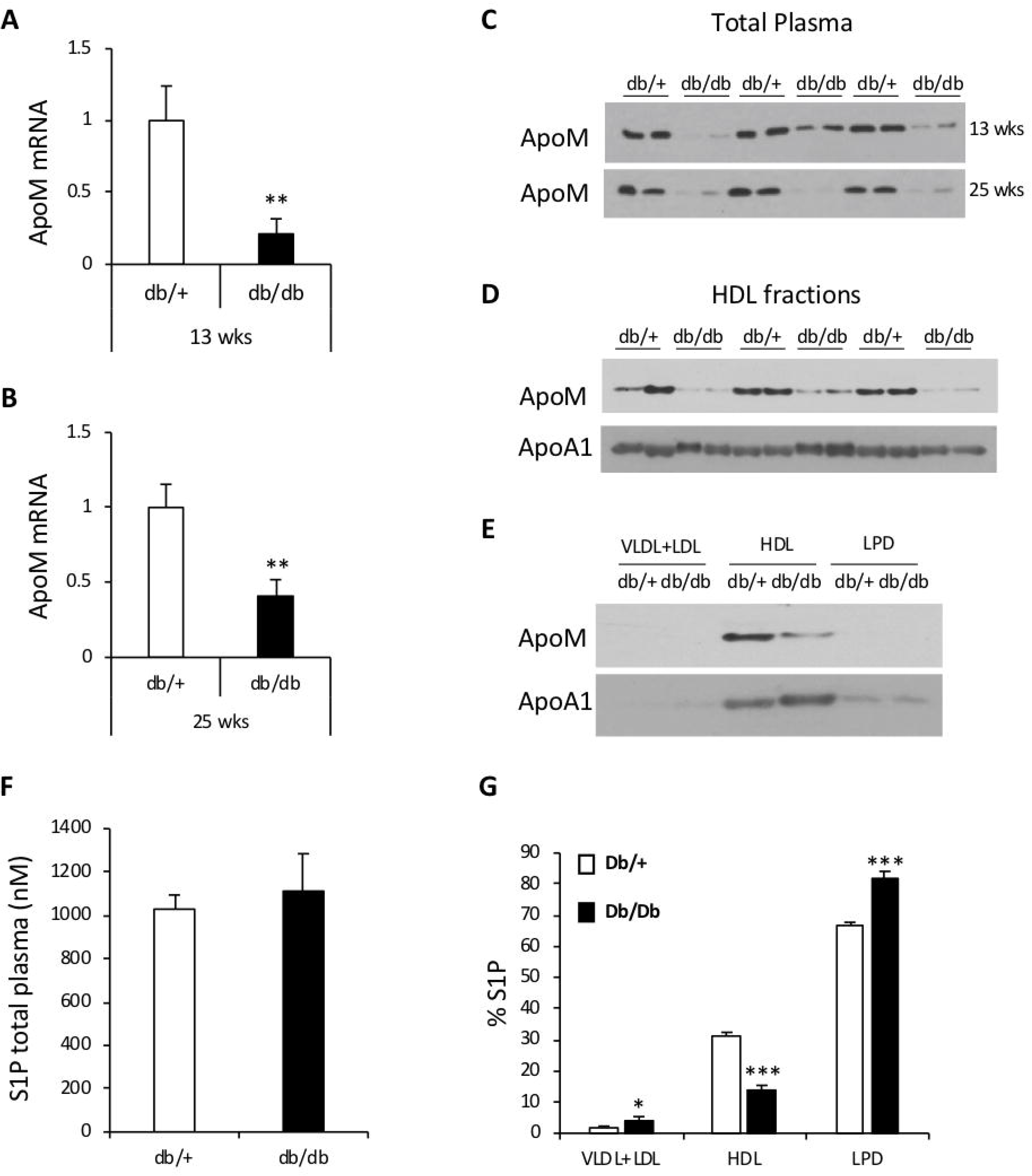
ApoM and S1P in db/db mice. **A)** Liver *Apom* gene expression in 13 week-old db/db mice and db/+ controls (n=6/group). **B)** Liver *Apom* gene expression in 25 week-old mice (n=6/group). **C)** Western blot of ApoM in total plasma from 13 week-old (top) and 25 week-old mice (bottom). **D)** Western blot of ApoM and ApoA1 in HDL fractionated by sequential density ultracentrifugation from plasma of 13 week-old mice. **E)** Representative western blot of ApoM and ApoA1 in ultracentrifuge-fractionated plasma from 13 week-old mice. **F)** Total plasma S1P levels. **G)** Distribution of S1P in plasma fractions. LPD=lipoprotein depleted. Data are presented as mean ± SEM. *P<0.05, **P<0.01, ***P<0.001 by student’s t-tests.

We had originally expected that FoxOs would be constitutively activated in db/db mice, so we investigated whether the expression of other known FoxO targets are affected in these mice. We observed that *G6Pc* was not elevated in our db/db mice at either age, although *Igfbp1* was (Supplementary Fig. 1d). *Gck* is normally suppressed by FoxOs^5, 12^, but it was increased in 13-week-old db/db mice (Supplementary Fig. 1d). Therefore, based on these canonical FoxO targets in the glucose metabolism pathway, there is no clear induction of FoxO targets that would indicate constitutive FoxO activation. On the other hand, we have previously reported that *Scarb1* and *Lipc*, two hepatic genes involved in HDL-cholesterol uptake into liver, are induced by FoxOs^14^. We found both *Scarb1* and *Lipc* are reduced in db/db mice (at 25 weeks and both ages, for *Scarb1* and *Lipc*, respectively) (Supplementary Fig. 1e). This suggests the possibility that in db/db mice, a subset of FoxO target genes related to lipoprotein metabolism are inactivated, through an unknown mechanism.

The second insulin resistance model that we studied were diet-induced obese mice. We fed C57BL/6J mice with a high fat diet starting at 6-weeks of age, until harvesting tissues when the mice were 13 or 29 weeks old. We observed an increase in the body weight, glucose, and insulin levels of the high fat-fed mice, as expected (Supplementary Fig. 2a-c). But we did not see an effect on the *Apom* mRNA expression or at the protein level (Supplementary Fig. 2d-f). Altogether, these data suggest that ApoM expression and S1P binding to HDL are affected in some, but not all, models of hyperinsulinemia and insulin resistance.

## DISCUSSION

Our findings show that FoxOs promote hepatic ApoM expression, and the FoxO-ApoM pathway is required for S1P to associate with HDL. Moreover, our data indicate that the majority of plasma ApoM arises from hepatic secretion, and that hepatic ApoM is required only for HDL-associated S1P, not total plasma S1P. We recently demostrated that hepatic FoxOs also promote clearance of HDL-cholesterol, by promoting expression of *Scarb1*, encoding scavenger receptor BI (SR-BI), and *Lipc*, encoding hepatic lipase^14^. Together with the novel data in this manuscript, these findings suggest that there are at least two independent mechanisms by which hepatic FoxOs regulate HDL composition.

S1P is a signaling molecule, and it is been reported that S1P can induce differential effects, depending on its chaperone, either ApoM or albumin^16, 20–23^. The molecular mechanisms for the differential effects between ApoM- and albumin-bound S1P remain under investigation. One possibility is the accessibility of S1P. Within the structure of ApoM, S1P interacts specifically with an amphiphilic pocket in the lipocalin fold^16^. Albumin serves as a promiscuous binding protein for various hydrophobic molecules, but it is unknown specifically how S1P binds to albumin. This raises the possibility that being bound to ApoM allows S1P to interact more efficiently with S1P receptors, although the binding of ApoM and S1P is strong, so the spontaneous release of S1P from ApoM is unlikely^39^. Another possibility invokes the preferential binding to HDL to certain cell types. SR-BI is a cell-surface receptor that binds HDL, and it has been shown to interact with S1PRs and allow activation by HDL-bound S1P^40^.

What is the consequence of FoxO inducing apoM and HDL-associated S1P? One possibility we considered was that this would play a role in FoxOs’ regulation of glucose homeostasis. This was suggested by data showing that several-fold overexpression of ApoM in wild-type mice improves glucose tolerance, and that ApoM-containing lipoproteins can stimulate insulin secretion from a mouse insulinoma cell line^35^. While L-FoxO1,3,4 mice *per se* have improved glucose tolerance, the rescue of ApoM in our experiment had no effect on glucose tolerance.

On the other hand, it is possible that decreased hepatic expression of ApoM may have other local and systemic consequences. For example, multiple publications have suggested a role for apoM-S1P in endothelial function, including increased phosphorylation of eNOS, decreased expression of immune cell adhesion molecules, and increased endothelial barrier function^16, 22–24^. Thus, it is possible that the low ApoM-S1P in L-FoxO1,3,4 mice cause them to have impairments in endothelial functions.

It is of interest to compare L-FoxO1,3,4 mice to *Apom*^−/−^ mice. L-FoxO1,3,4 mice have a near-total absence (∼90% reduction) of hepatic *Apom* mRNA and protein, and a similarly potent reduction of plasma ApoM. This causes a substantial reduction in HDL-associated S1P. However, the S1P bound to albumin is compensatorily increased, such that L-FoxO1,3,4 mice have no differences in total plasma S1P levels compared to the control mice. These findings indicate that: (i) most ApoM in plasma arises from hepatocytes and (ii) hepatic ApoM is either not required for maintaining total plasma S1P content, or very low levels of ApoM are sufficient. In contrast, whole-body *Apom*^−/−^ mice have a reduction in total plasma S1P by 46% compared to wild-type mice^16^. The contrast between these two mouse models suggests the possibility that other, non-hepatocyte cells that express ApoM are involved in maintaining total plasma S1P levels.

FoxOs are inactivated by insulin, and it is widely believed that they are constitutively active in settings of insulin resistance^36^. Interestingly, in insulin resistant db/db mice, where FoxOs may be expected to be constitutively active, ApoM is reduced. Xu et al. previously showed that hepatic *Apom* expression and the levels of ApoM in plasma are significantly lower in db/db mice^41^. We confirmed these findings in multiple cohorts of db/db mice, at different ages (12-13 weeks and 25 weeks), and extended them by showing that db/db mice have reductions in the levels of S1P bound to HDL, with no differences in total plasma S1P levels. Consistent with the low ApoM, we also found that other FoxO targets involved in HDL homeostasis, *Scarb1* and *Lipc*^14^ are also reduced in db/db mice. This is in contrast to other canonical FoxO targets in the glucose metabolism pathway, whose expression are varyingly suggestive of FoxO being active, inactive, or unaffected in these db/db mice. Thus, the molecular basis underlying the decrease specifically in lipoprotein-related FoxO targets remains to be elucidated. Potential explanations may be suggested by comparing the db/db mice with the diet induced obese mice, which had no differences in *Apom* expression. In the db/db mice the levels of glucose and insulin are increased ∼2-fold and ∼20-30-fold, respectively. These changes were milder in the diet induced obese mice, where glucose increased 1.4-fold and insulin increased 6-8-fold. Perhaps the extreme hyperinsulinemia in db/db mice is capable of inactivating a subset of FoxOs. Another possibility is that FoxOs’ effects on ApoM expression could be sensitive to glucose levels or other hormones. FoxOs are regulated by multiple different posttranslational modifications in addition to the effects of insulin signaling to suppress FoxO activity via Akt-mediated phosphorylation and nuclear exclusion^42, 43^. In response to hyperglycemia and oxidative stress, FoxOs are acetylated, which causes accelerated FoxO degradation^44^. In response to glucagon and downstream calcium signaling, FoxOs are phosphorylated on alternative phosphorylation sites that cause its activation^45, 46^. Thus, in certain settings of murine insulin resistance, a subset of FoxOs may be inactivated, causing decreased expression of *Apom* and potentially other FoxO targets.

Individuals with type 2 diabetes have low levels of high-density lipoprotein (HDL)-cholesterol, and these low HDL-cholesterol levels are inversely correlated to cardiovascular disease. However, clinical trials have demonstrated that raising HDL-cholesterol *per se* is generally insufficient to reduce coronary disease^47^. It is possible that other aspects of HDL are defective in diabetes, contributing to cardiovascular risk. A recent paper has shown that diabetes patients have reduced S1P-HDL levels and impaired HDL cardioprotective function^48^. Moreover, plasma ApoM and S1P levels are inversely associated with mortality in diabetes patients^49^. These findings suggest that an alternative approach to lowering cardiovascular risk in diabetes would be to increase ApoM and S1P, which may have endothelium-protecting effects^16, 22–24^.

## METHODS

### Mice and diets

All experiments were approved by the Columbia University Institutional Animal Care and Use Committee. L-FoxO1,3,4 mice have been previously described^5, 9^. Males at least 12 weeks old were studied, except in studies of 2-day-old pups, which were of mixed sex. For the acute FoxO depletion experiments, adult *Foxo1^fl/fl^*, *Foxo3^fl/fl^*, and *Foxo4^fl/Y^* control mice were transduced with adeno-associated virus (serotype 8) expressing Cre recombinase driven by the hepatocyte-specific Tbg promoter (AAV8.Tbg. Cre) or control virus (AAV.GFP). Mice were injected intravenously with 1×10^11^ virus particles/mouse, 4 weeks prior to euthanasia. AAV8.Tbg.Cre was a gift of Morris Birnbaum (Perelman School of Medicine, University of Pennsylvania, Philadelphia, Pennsylvania USA). For the adenovirus experiments, adult male mice were injected intravenously with murine ApoM adenovirus (Welgen) 0.5 × 10^9^ virus particles/gram of body weight, 8 days prior to euthanasia. For the db/db studies, male db/db and db/+ mice were purchased from Jackson Laboratory and were studied when they were 12, 13 or 25 weeks old. For the diet induced obesity studies, male C57BL/6J mice were purchased from Jackson Laboratory when they were 6 weeks old. Mice were fed either a standard chow diet (Purina) or a high fat diet containing 60 kcal% from fat (ResearchDiets D12492) until they were 13 or 29 weeks old. Western type diet, containing 42% kcal from fat and 0.2% cholesterol, is from Harlan Teklad (TD.88137). Mice were maintained on a 12-hour light/12-hour dark cycle, with the dark cycle occurring between 7:00 pm and 7:00 am.

### Primary hepatocytes studies

Primary hepatocytes were isolated from male mice via collagenase perfusion, as previously described^12^. We plated them on collagen-coated culture ware for 2 hours. Following attachment, hepatocytes were transduced with FoxO1-ADA (T24A-S253D,S316A mutations)^27^, FoxO1-ADA-DBD T24A, S253D and S316A mutations (ADA) + N208A and H212R mutations (DBD)) ^28^ or FoxO1-Δ256 (AA1-256, truncated form)^34^ for 16 hours.

### mRNA and protein expression

Liver and hepatocyte RNA was extracted using TRIzol (Invitrogen, Thermo Fisher Scientific). cDNA was obtained using High-Capacity cDNA Reverse Transcription Kit (Applied Biosystems). Quantitative PCR was performed with iTaq Universal SYBR Green Supermix (Bio-Rad). 36b4 was used as housekeeping gene for normalization. Primer sequences available upon request. Western blots used the primary antibodies directed against the following proteins: ApoM (LSBio), ApoA1 (Meridian Life Science), β-actin (Cell Signaling).

### Metabolic tests

Blood glucose was measured using Breeze2 monitor and strips (Bayer). Insulin ELISAs were from Millipore. For intraperitoneal glucose tolerance tests, mice were fasted for 16 hours and injected intraperitoneally with glucose (2 g/kg). We obtained blood samples at 0, 15, 30, 60, 90 and 120 min after the injection and measured glucose levels.

### Plasma lipoprotein analysis

VLDL+LDL (d<1.063 g/mL), HDL (1.063<d<1.210 g/mL) and lipoprotein depleted (d>1.210 g/mL) fractions were separated by sequential density ultracentrifugation. 70 μl of total plasma was separated by using the Optima MAX-TL Ultracentrifuge with TLA-100 rotor (Beckman Coulter). Plasma lipoproteins were also analyzed by running 200 μl of plasma onto a fast protein liquid chromatography system consisting of Superose6 10/300 GL column (Amersham Pharmacia Biotech), and fractions were collected using the fraction collector FC-204 (Gilson). Plasma cholesterol was measured using a colorimetric assay (Wako).

### S1P quantification

Bioactive sphingolipids were extracted from total plasma or VLDL+LDL, HDL and lipoprotein depleted fractions, run on liquid chromatography-tandem mass spectrometry (LC-MS/MS), and quantitated using stable isotope-labeled internal standards.

### Statistics

Statistical analysis was performed using SPSS 11.0 statistical software. Data are expressed as the mean ± SEM. Significance at the *P*<0.05 level was assessed by Student’s *t*-test.

## ACKNOWLEGEMENTS

The authors would like to gratefully acknowledge Marit Westerterp (University of Groningen), Alan Tall (Columbia University), and Domenico Accili (Columbia University) for discussions and suggestions. We also acknowledge technical assistance from K. Zhong, A. Flete, T. Kolar (all from Columbia University). This work was funded by NIH R01HL125649 (to RAH), ADA 1-17-PMF-017 (to MCI). We also acknowledge the Irving Clinical Translational Research Center, funded by NIH UL1 TR001873.

---

**Supplementary Figure 1.**
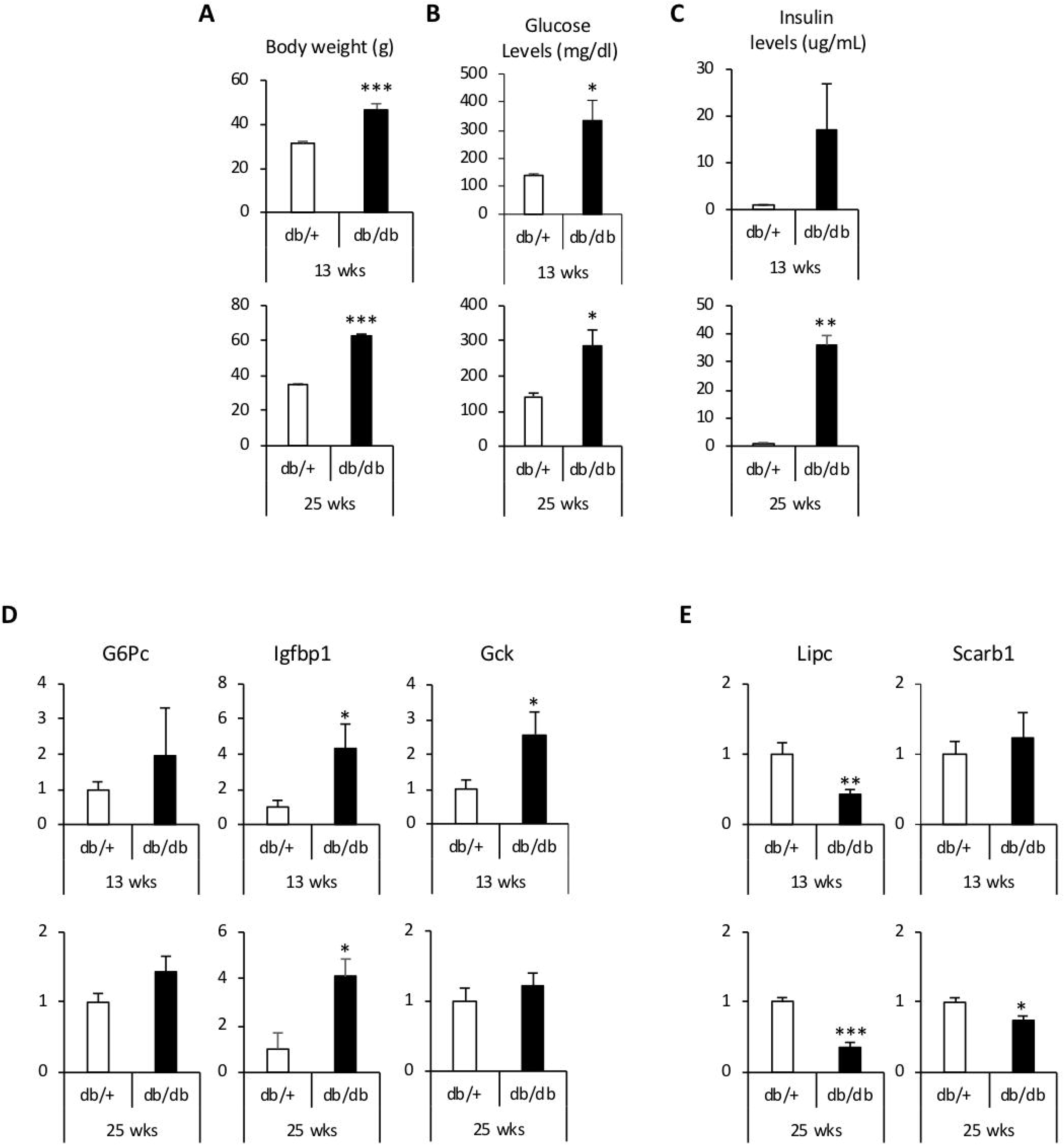
Metabolic parameters and hepatic gene expression in db/db mice. Top: 13 week-old mice, Bottom: 25 week-old mice. **A)** Total body weight. **B)** Plasma glucose levels after 5 hours fasting. **C)** Insulin levels after 5 hours fasting. **D)** *G6pc, Gck* and *Igfbp1* gene expression in liver **E)** *Lipc* and *Scarb1* gene expression in liver. (n=6/group for all panels) Data are presented as mean ± SEM. *P<0.05, **P<0.01, ***P<0.001 by student’s t-tests.

**Supplementary Figure 2.**
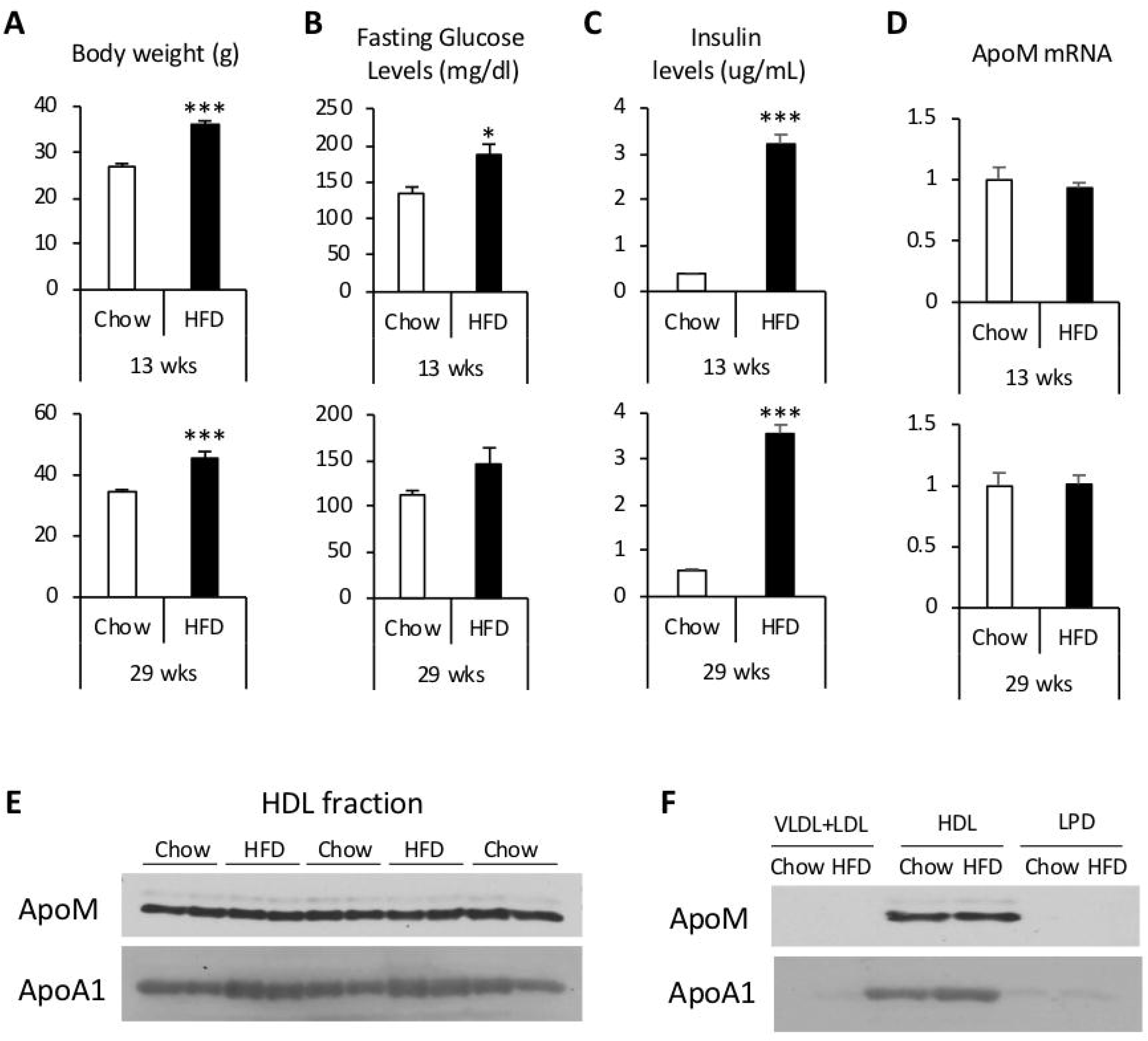
Metabolic parameters, hepatic gene expression, and HDL-ApoM in diet induced obese mice. Top: 13 week-old mice, Bottom: 29 week-old mice. HFD=high fat diet. **A)** Total body weight. **B)** Plasma glucose levels after 5 hours fasting. **C)** Insulin levels after 5 hours fasting. **D)** *Apom* gene expression in liver. **E)** Western blot of ApoM and ApoA1 in HDL fractionated by sequential density ultracentrifugation from plasma of 13 week-old mice. **F)** Representative western blot of ApoM and ApoA1 in ultracentrifuge-fractionated plasma from 13 week-old mice. (n=6/group for all panels) Data are presented as mean± SEM. *P<0.05, **P<0.01, ***P<0.001 by student’s t-tests.

